# Canonical autophagy remains inactive in induced pluripotent stem cells and neuronal progenitor cells following DNA damage induced by BPDE or etoposide

**DOI:** 10.1101/2025.05.22.655294

**Authors:** Seda Akgün, Thomas Lenz, Annika Zink, Karina Stephanie Krings, Sebastian Wesselborg, María José Mendiburo, Alessandro Prigione, Kai Stühler, Björn Stork

## Abstract

(Macro-)Autophagy is a key cellular stress response mediating the recycling of long-lived or damaged proteins and organelles. In stem cells, autophagy is essential for the decision between quiescence, self-renewal and differentiation. We observed that induced pluripotent stem cells (iPSCs) and thereof derived neural progenitor cells (NPCs) have a functional autophagy machinery, as shown by starvation-induced autophagic flux and ULK1 activation. Using the human iPSC line iPS11 and thereof derived NPCs (niPS11), we investigated whether genotoxic stress induced by benzo[a]pyrene diolepoxide (BPDE) or etoposide can similarly activate autophagy, as previously reported for cancer cell lines. While both BPDE and etoposide induced the DNA damage markers phospho-p53 Ser15 and γH2AX and slightly altered the expression of DNA repair proteins such as XPC, they did not trigger autophagic flux in either iPSCs or NPCs. After genotoxin treatment, ULK1 activation was only observed in NPCs, but this was not sufficient to trigger a significant downstream autophagic response. Mass spectrometry revealed minimal proteomic changes in iPSCs and moderate changes in NPCs, mainly involving mitotic regulators. These results suggest that genotoxic agents do not strongly affect canonical autophagy in pluripotent stem cells or their neural derivatives despite an otherwise responsive autophagic system.

## INTRODUCTION

(Macro-)autophagy represents an intracellular stress response responsible for the recycling of long-lived or damaged proteins and organelles. During this process, the cargo to be degraded becomes engulfed within double-membraned vesicles termed autophagosomes. The outer membranes of these autophagosomes fuse with lysosomes, forming autolysosomes. Within lysosomes, lysosomal hydrolases degrade the cargo, and the resulting building blocks such as amino or fatty acids are transferred back to the cytosol where they are available again for ATP production, protein synthesis, etc. Autophagy occurs at basal levels in most cell types and ensures cellular homeostasis. However, an autophagic response can also be induced upon stress conditions such as nutrient deprivation, protein aggregation, infection with intracellular pathogens, or DNA damage. Autophagy is executed by autophagy-related (ATG) proteins and non-ATGs, mediating all steps of the autophagy pathway, i.e., initiation, nucleation of the phagophore, expansion of the autophagosomal membrane, maturation of autophagosomes, and fusion with lysosomes. The initiation of autophagy is centrally regulated by the autophagy-inducing ULK1 complex, containing the Ser/Thr kinase unc51-like kinase 1 (ULK1) and the associated factors ATG13, ATG101, and FIP200 (Yamamoto et al., 2023). Two frequently used autophagy marker proteins are the microtubule-associated proteins 1A/1B light chain 3 (MAP1LC3, in brief LC3) and sequestosome-1 (SQSTM1)/p62, which function in autophagosome biogenesis and cargo recruitment (Mizushima & Murphy, 2020).

Autophagy plays a particularly important role in stem cell populations, as they are dependent on intracellular quality control and the maintenance of cellular homeostasis. Autophagic processes have been studied in various stem cell types, including embryonic stem cells, various tissue stem cells (e.g. hematopoietic or neural stem/progenitor cells, NSPCs), cancer stem cells and induced pluripotent stem cells (iPSCs) (Rodolfo et al., 2016). Previous research suggests that autophagy plays a central role in the decision between quiescence, self-renewal and differentiation (Rodolfo et al., 2016). In NSPCs, cytoprotective autophagy is involved in both maintenance and neuronal differentiation (Casares-Crespo et al., 2018). It has been shown that FIP200, a component of the autophagy-inducing ULK1 kinase complex, is essential for these two functions, especially in the postnatal brain (Wang et al., 2013). On the other hand, it could be shown in the mouse model that inhibition of autophagy reduces the irradiation-induced loss of NSPCs (Wang et al., 2017).

The DNA damage response (DDR) is a cellular stress response usually initiated upon genotoxic stress. Generally, a DDR is initiated with the detection of the DNA lesion and the recruitment of factors mediating DNA repair. DNA repair in turn can be executed by five different pathways, depending on the type of DNA lesion. These pathways are 1) base excision repair (BER), 2) nucleotide excision repair (NER), 3) mismatch repair (MMR), 4) homologous recombination (HR) and 5) non-homologous end joining repair (NHEJ) (Sadoughi et al., 2021). In vertebrate cells, the DDR is controlled by three related kinases: ataxia telangiectasia mutated (ATM), ataxia telangiectasia and Rad3 related (ATR), and DNA-dependent protein kinase (DNA-PK) (Blackford & Jackson, 2017). On the level of these three kinases, the crosstalk between the DDR and autophagy is initiated. It is generally assumed that autophagy provides the metabolic resources to enable DNA repair. On the molecular levels, it has been demonstrated that ATM, ATR and DNA-PK regulate autophagy signaling via transcriptional or post-translational control (Juretschke & Beli, 2021). The transcriptional control might be executed via the activation of p53 or TFEB, or the nuclear exclusion of FOXK (Juretschke & Beli, 2021). The posttranslational control mainly involves the inactivation of the mammalian target of rapamycin (mTOR) and/or the activation of the AMP-activated protein kinase (AMPK) (Juretschke & Beli, 2021), two key upstream regulators of the above-described autophagy-inducing ULK1 complex. In turn, autophagy modulates DNA repair pathways (Gomes et al., 2017). Finally, it has recently been demonstrated that autophagy exerts a direct role in the repair of DNA lesions, via TEX264-mediated selective autophagy of topoisomerase 1 cleavage complexes (TOP1cc) DNA lesions (Lascaux et al., 2024).

Autophagy induction by anticancer drugs has been reported for several cell lines, but the effects in stem cells are largely unknown. In this study, we aimed at investigating how genotoxins affect autophagy signaling in iPSCs and thereof differentiated NPCs. We utilized benzo[a]pyrene diolepoxide (BPDE) and etoposide as model compounds. BPDE is a metabolite of benzo[a]pyrene (BaP), a polycyclic aromatic hydrocarbon (PAH) found in tobacco smoke, smog and other combustion products, and it forms adducts with nitrogen-containing bases of the DNA (Zhao et al., 2024). Etoposide in turn is a potent topoisomerase II poison, causing double-stranded DNA breaks (Bailly, 2023). We found that neither compound elicited a significant autophagic response in iPSCs and NPCs, albeit the autophagic machinery is both present and functional in these cells.

## RESULTS

### Characterization of iPSCs and NPCs

In this study, we made use of the iPSC lines iPS11 (derived from human foreskin fibroblasts, Alstem) and iPS12 (human mesenchymal stromal cells, Alstem). Both iPSC lines ectopically express OCT4, SOX2, KLF4, and L-MYC. OCT4 expression was confirmed by immunoblotting (Figure 1A) and by immunofluorescence microscopy (Figure 1B). Differentiation of iPSCs to neural progenitor cells (NPCs) was done as previously described (Zink et al., 2021) and as depicted in Figure 1C, and the resulting cell line was designated niPS11. Expression of NPC marker proteins Pax6 and Nestin was confirmed in niPS11 by immunoblotting and immunofluorescence, respectively (Figure 1A and 1B).

**Figure 1:**
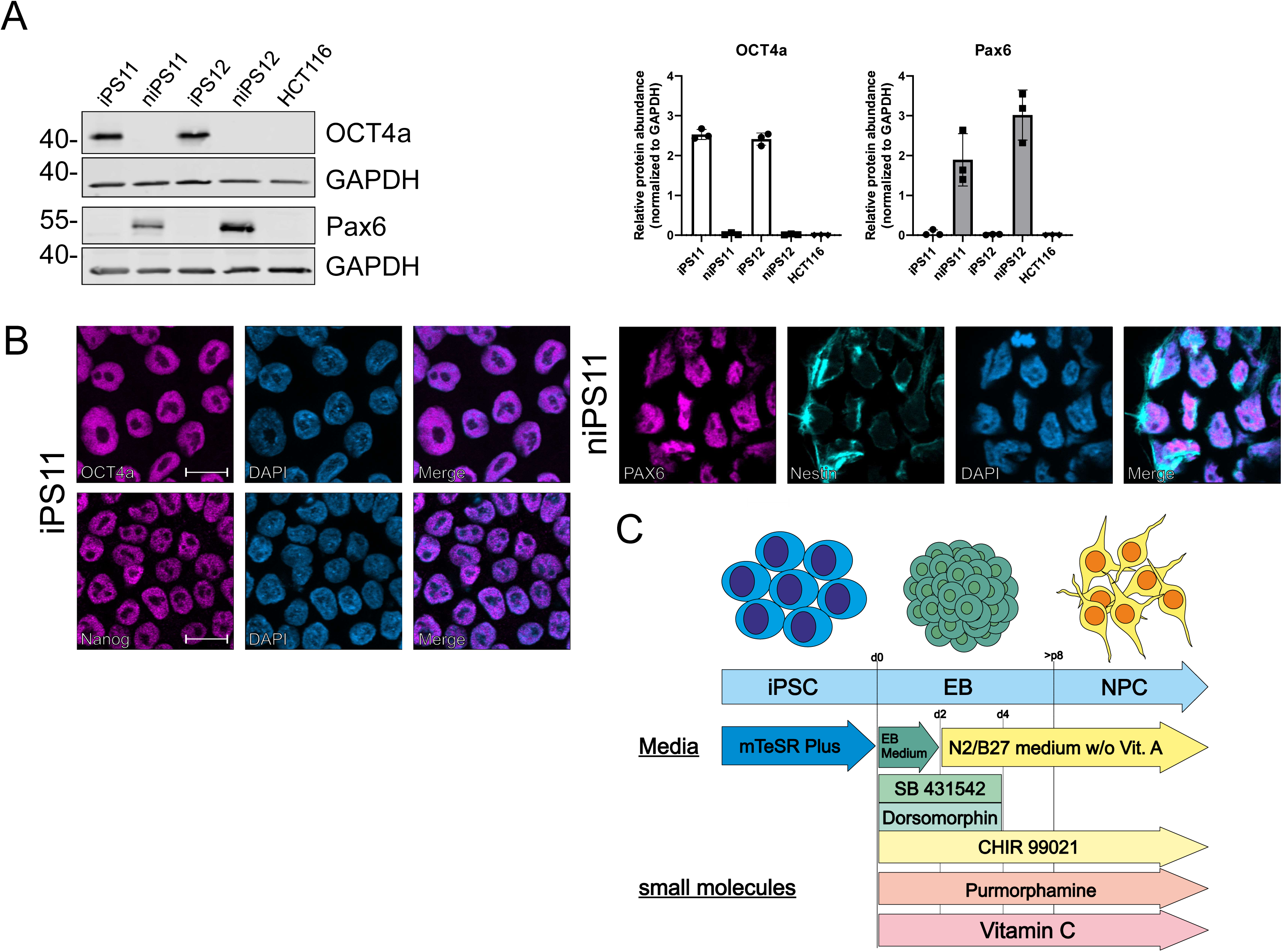
Verification of induced pluripotent stem cells and thereof differentiated neural progenitor cells. (**A**) Induced pluripotent stem cells, neural progenitor cells and HCT116 were lysed, and cellular lysates were immunoblotted for OCT4a, Pax6 and GAPDH respectively. One representative immunoblot is shown. The quantifications of indicated ratios are from three independent experiments (means ± SD). (**B**) Cells were fixed and stained for pluripotency markers OCT4a and Nanog in iPS11 and neural markers Pax6 and Nestin in niPS11 and visualized by immunofluorescence. Scale bar: 10 µm. (**C**) Differentiation of iPSCs into NPCs was achieved by SMAD and AMPK inhibition via SB 431542 and Dorsomorphin, directed and sustained as neural progenitor cells by supplementation of above mentioned compounds.

### iPSCs and NPCs reveal starvation-inducible autophagic capacity

In order to evaluate genotoxin-induced autophagy in iPSCs and thereof differentiated NPCs, we first investigated the general starvation-inducible autophagic capacity of these two cell models. For that, we starved the cells in the absence or presence of bafilomycin A_1_, which is a vacuolar-type H^+^-translocating ATPase (V-ATPase) inhibitor blocking autolysosomal degradation, and analyzed autophagy by immunoblotting for the autophagy markers phospho-ULK1 (Ser758), LC3, and p62/SQSTM1. The autophagy-inducing kinase ULK1 is phosphorylated at Ser758 by mTOR complex 1 (mTORC1) and thus kept in an inhibited state. Dephosphorylation of this site correlates to the induction of autophagy (Dorsey et al., 2009; Kim et al., 2011; Shang et al., 2011). Lipidated LC3 (LC3-II) decorates the inner and outer surfaces of autophagic membranes and recruits both components of the autophagic machinery and cargo to be degraded. P62 is an autophagy receptor mediating the recruitment of autophagic cargo. In both cell lines, starvation induced autophagic flux as determined by these three markers, i.e., reduced phosphorylation of ULK1 at Ser758, increased LC3-II turnover (difference in LC3-II levels with and without bafilomycin A_1_), and increased p62 turnover (difference in p62 levels with and without bafilomycin A_1_) (Figure 2A and 2B). Collectively, these data indicate the “general” autophagic capacity of both iPSCs and NPCs.

**Figure 2:**
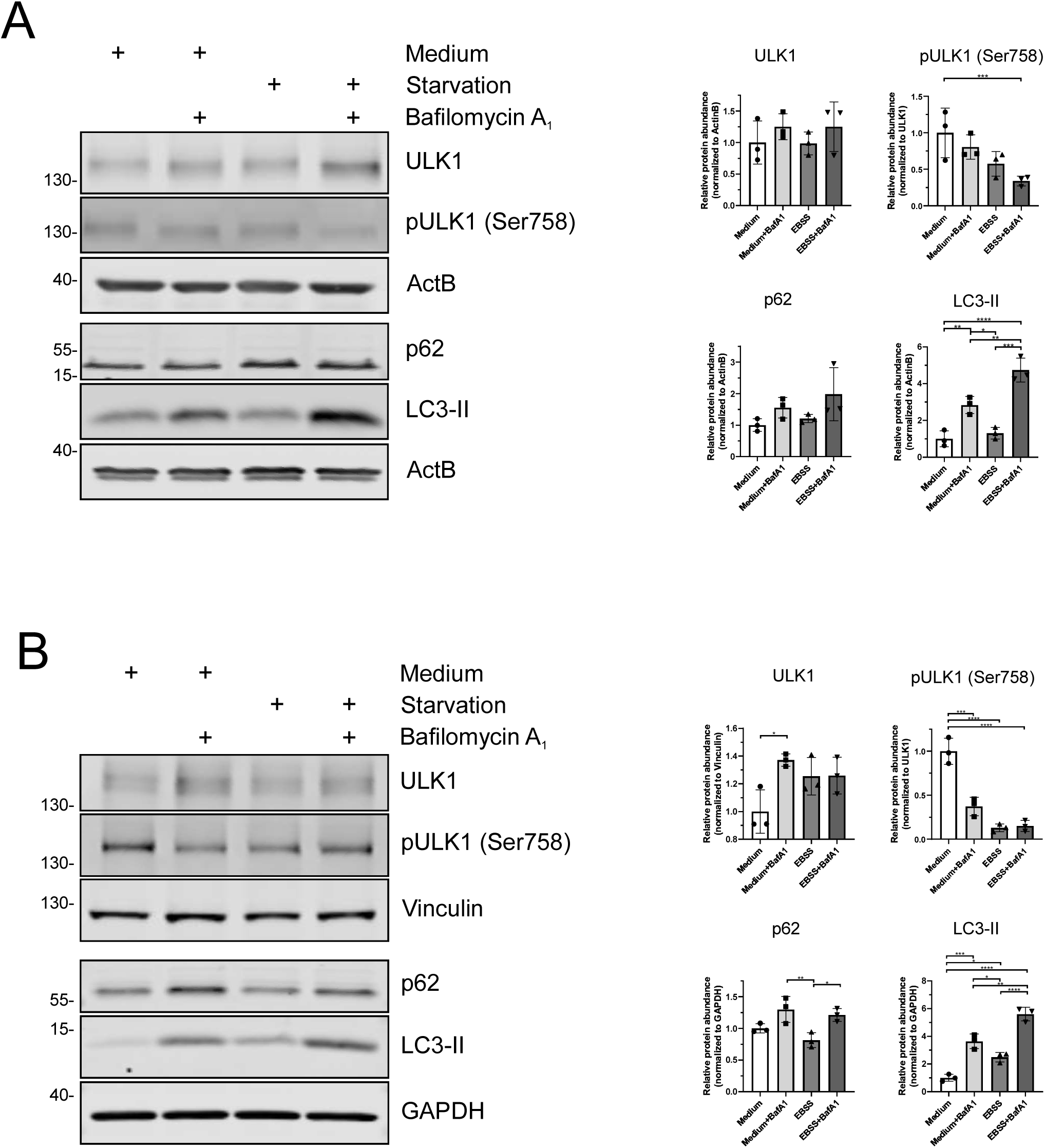
Both stem cell types show canonical autophagic capacity. (**A & B**) Both cell types were treated with either cultivation medium or serum- and amino acid-free EBSS for 2 h. For the accumulation of the lysosome-associated proteins LC3-II and p62, V-ATPase inhibitor bafilomycin A_1_ was additionally supplemented to each medium. Afterwards, cells were lysed, and cellular lysates were immunoblotted for ULK1, phospho-ULK1 Ser758, Vinculin, SQSTM1/p62, LC3 and GAPDH. One representative immunoblot is shown. Results show mean + SD of three independent experiments. For statistical analysis ordinary one-way ANOVA (Tukeýs multiple comparisons test) was utilized to compare means of genotoxin-treated samples to DMSO. * p < 0.05, ** p < 0.01, *** p < 0.001, **** p < 0.0001.

### BPDE and etoposide induce DNA damage in iPSCs and NPCs

In order to determine suitable concentrations of the genotoxins BPDE and etoposide for our autophagy assays, we first determined IC_20_ values. For that, we performed MTT assays in both cell models and analyzed viability after 24, 48 and 72 hours, respectively (Suppl. Figure 1). As treatment scheme, cells were exposed to etoposide for 24 h and subsequent medium exchanges were without etoposide. In contrast, BPDE was freshly supplemented to the cells every 24 hours. As control, we used the colon carcinoma cell line HCT116. For both genotoxins, IC_20_ values were lower in iPS11 as compared to niPS11 over all time points, indicating a higher sensitivity of iPS11 for DNA-damaging agents (Suppl. Figure 1). Next, we assessed DNA damage by immunoblotting for phospho-p53 (Ser15) and phospho-H2AX (Ser139; γH2AX). Ser15 of p53 can become phosphorylated by ATM, ATR, and DNA-PK, and this phosphorylation prevents p53 from associating with MDM2, ultimately leading to the accumulation and activation of p53 following DNA damage (Shieh et al., 1997; Tibbetts et al., 1999). Similarly, phosphorylation of H2AX at Ser139 is also mediated by the mentioned kinases upon DNA damage (Burma et al., 2001; Rogakou et al., 1999). In iPS11, both phospho-substrates were detectable upon both treatments, with a faster kinetic upon etoposide treatment compared to BPDE treatment (Figure 3A). Of note, γH2AX was also clearly induced by starvation (EBSS). This was also the case for niPS11 and HCT116 (Figures 3B and 3C), and this observation might be attributed to a p38-dependent phosphorylation of H2AX (Lu et al., 2008). In niPS11, the kinetics of both treatments are rather similar. In this model, γH2AX appeared to be sensitive to bafilomycin A_1_ treatment (Figure 3B). Generally, the DNA damage-induced phosphorylation of p53 and H2AX was clearly observable in HCT116, and also more distinct in comparison to untreated controls (Figure 3C).

**Figure 3:**
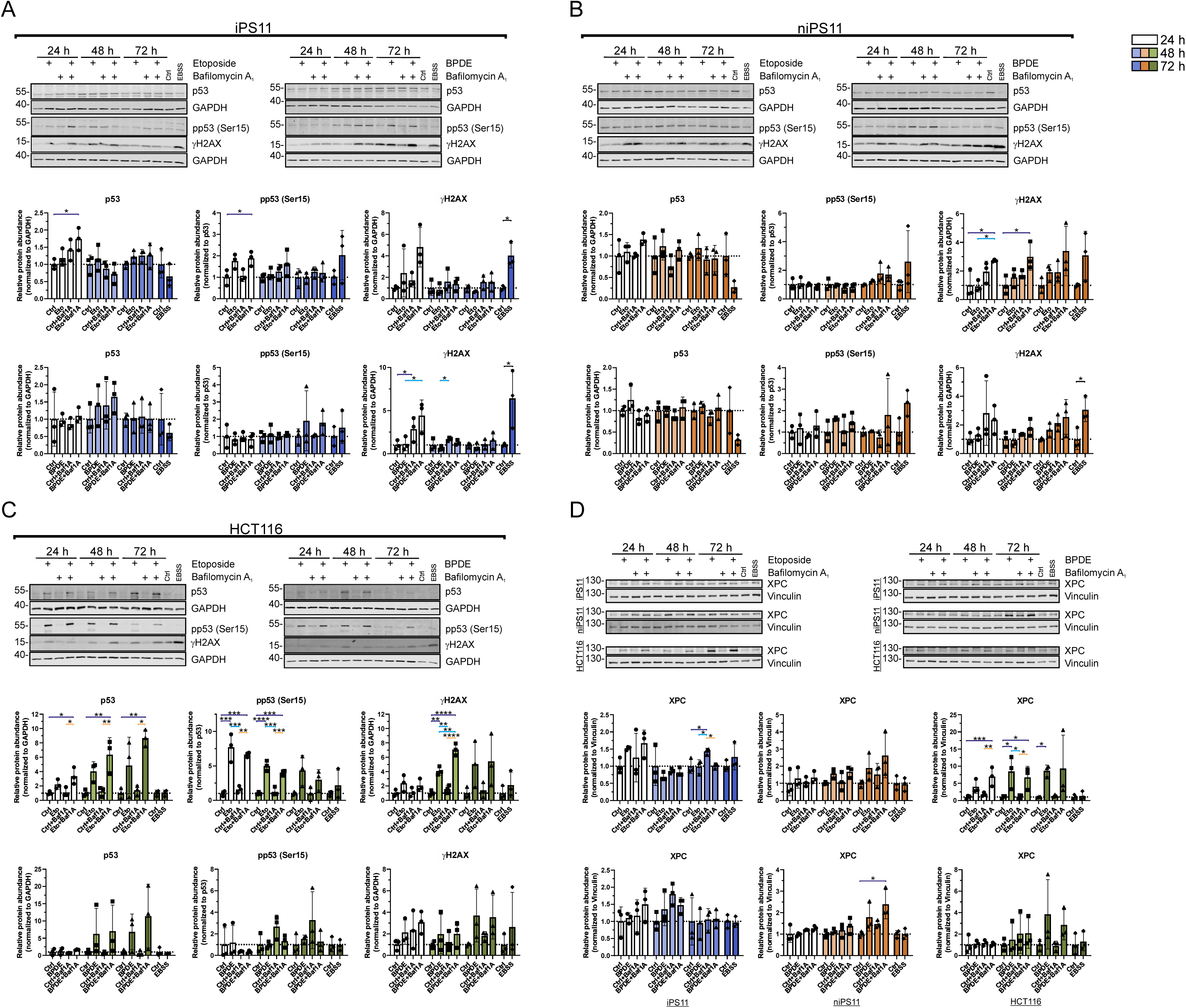
Stem cells show a more moderate impact on DNA damage marker compared to HCT116. (**A-D**) General DNA damage response proteins were investigated by immunoblotting. (**A**) iPS11, (**B**) niPS11 and (**C**) HCT116 were treated with corresponding IC_20_ dose for 24, 48 and 72 h. Therefore, etoposide was supplemented for 24 h and exchanged with genotoxin-free medium for 48 h and 72 h while BPDE was supplemented daily. 4 hours before harvesting medium was exchanged and 40 nM bafilomycin A_1_ was supplemented to labeled samples 2 h before lysis. Cells were lysed, and cellular lysates were immunoblotted for p53, phospho-p53 Ser15, phospho-H2AX Ser139 and GAPDH. (D) Furthermore, NER protein XPC was also immunoblotted as described above. One representative immunoblot is shown. Results show mean + SD of three independent experiments. For statistical analysis ordinary one-way ANOVA (Tukeýs multiple comparisons test) and student’s t-test were utilized to compare means of genotoxin-treated samples to DMSO. Significance bars are highlighted in different colors accordingly to involved condition: dark blue: control, light blue: genotoxin; orange: control + bafilomycin A_1_ and genotoxin + bafilomycin A_1_. * p < 0.05, ** p < 0.01, *** p < 0.001, **** p < 0.0001.

We next analyzed Xeroderma pigmentosum group C protein (XPC) levels. XPC generally functions as an initiator of global genome nucleotide excision repair (GG-NER), which repairs lesions generated by BPDE (Piberger et al., 2018). Although DNA double strand breaks are the main type of damage induced by etoposide, NER proteins have also been linked to topoisomerase II inhibitors (Rocha et al., 2016). XPC levels were clearly increased by both genotoxins in control HCT116 cells. In iPS11 and niPS11 cells, effects on XPC levels were less prominent, with the exception of BPDE-treated niPS11 cells (Figure 3D).

Collectively, these data indicate that both genotoxins generally induce a DNA damage response in iPS11 and niPS11 cells.

### ULK1 activation status is not affected by BPDE or etoposide in iPSCs

Since the main goal of our project was to investigate genotoxin-induced autophagy in stem cells and thereof differentiated cells, we next investigated the activation status of the autophagy-inducing kinase ULK1. For that, we analyzed ULK1 phosphorylation at Ser758 and Ser638 (another mTOR/AMPK-dependent phospho-site) upon etoposide or BPDE treatment. In iPS11, no alterations of ULK1 activation status were observable, neither for etoposide nor for BPDE (Figures 4A and 5A), although responsiveness towards starvation could be confirmed (see also figure 2A). In contrast, ULK1 activation (i.e., dephosphorylation at Ser758 and Ser638) was detected in niPS11 (Figures 4B and 5B), although the extent of ULK1 activation was not as strong as with starvation. Of note, the effect of etoposide treatment was indifferent in HCT116, with rather decreased Ser758 phosphorylation and increased Ser638 phosphorylation (Figure 4C). With regard to BPDE, a consistent pattern of ULK1 activation was observed in HCT116 cells (Figure 5C). These data indicate that early autophagy events such as the activation of ULK1 do not occur in iPSCs upon genotoxin treatment, whereas this appears to be the case in NPCs.

**Figure 4:**
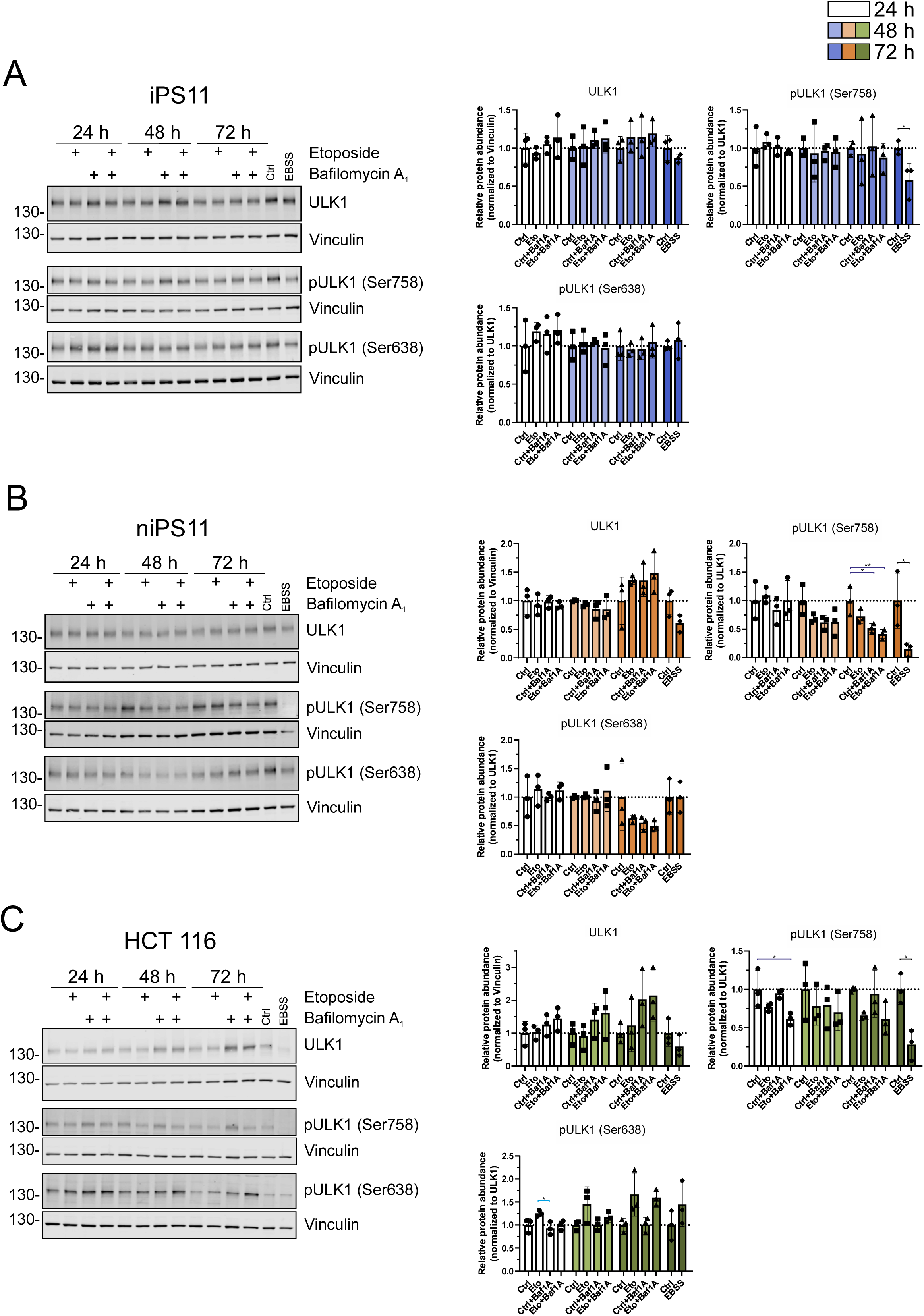
ULK1 activation status is not affected in iPS11 after etoposide treatment. (**A**) iPS11 (**B**) niPS11 and (**C**) HCT116 were treated with corresponding IC20 dose of etoposide and lysed after 24, 48, 72 h. Cellular lysates were immunoblotted for ULK1, phospho-ULK1 Ser758 and phospho-ULK1 Ser638 respectively. One representative immunoblot is shown. Results show mean + SD of three independent experiments. For statistical analysis ordinary one-way ANOVA (Tukeýs multiple comparisons test) and student’s t-test were utilized to compare means of genotoxin-treated samples to DMSO. Significance bars are highlighted in different colors accordingly to involved condition: dark blue: control, light blue: genotoxin; orange: control + bafilomycin A_1_ and genotoxin + bafilomycin A_1_. * p < 0.05, ** p < 0.01, *** p < 0.001, **** p < 0.0001.

**Figure 5:**
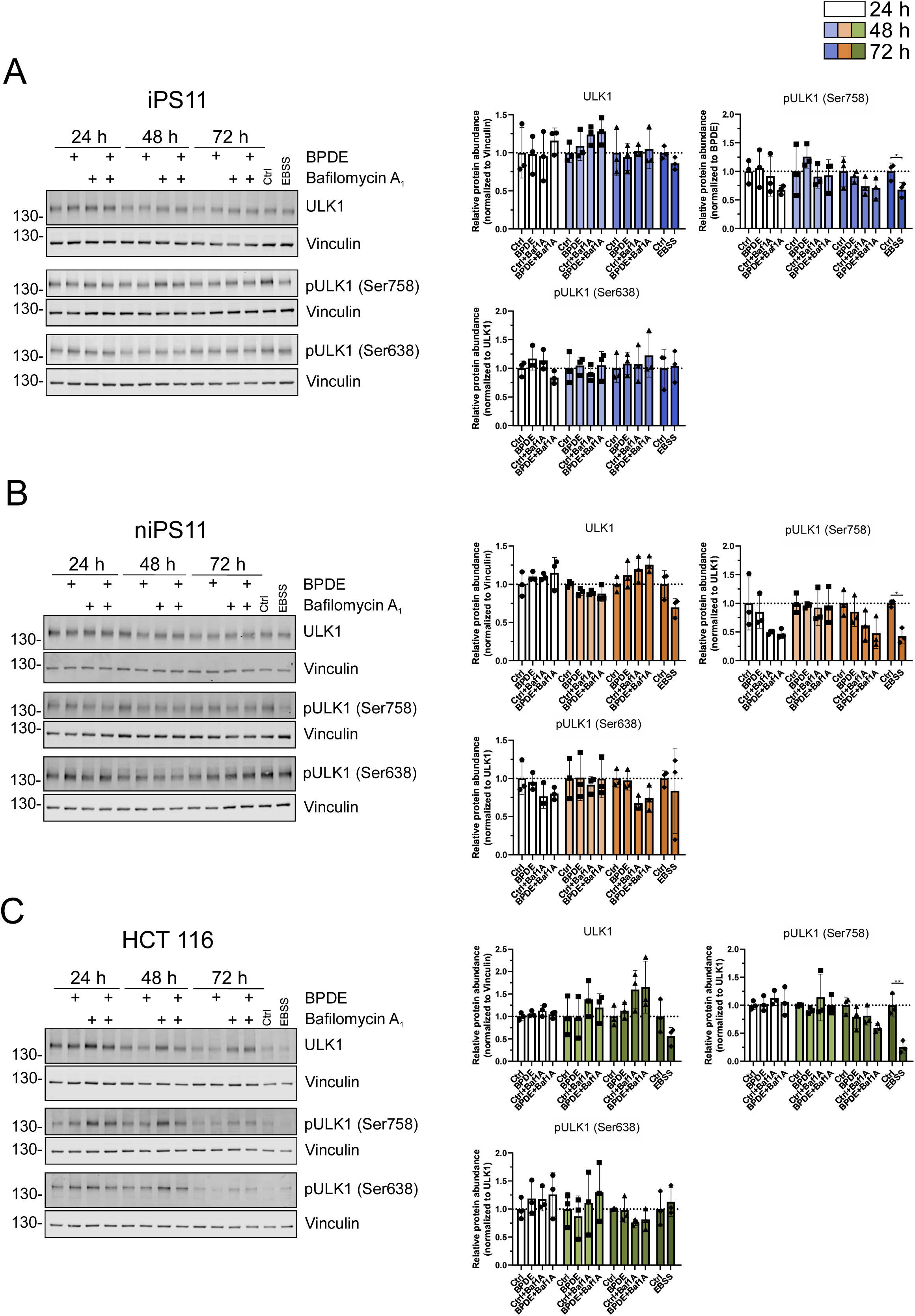
BPDE exposure does not influence ULK1 phosphorylation. (**A**) iPS11 (**B**) niPS11 and (**C**) HCT116 were treated with corresponding IC20 dose of BPDE and lyzed after 24, 48, 72 h. Cellular lysates were immunoblotted for ULK1, phospho-ULK1 Ser758 and phospho-ULK1 Ser638 respectively. One representative immunoblot is shown. Results show mean + SD of three independent experiments. For statistical analysis ordinary one-way ANOVA (Tukeýs multiple comparisons test) and student’s t-test were utilized to compare means of genotoxin-treated samples to DMSO. Significance bars are highlighted in different colors accordingly to involved condition: dark blue: control, light blue: genotoxin; orange: control + bafilomycin A_1_ and genotoxin + bafilomycin A_1_. * p < 0.05, ** p < 0.01, *** p < 0.001, **** p < 0.0001.

### BPDE or etoposide do not induce autophagic flux in iPSCs or NPCs

Next to early ULK1 activation, we also investigated the effect of the two genotoxins on autophagic flux. For that, we monitored again turnover of LC3 and p62. Neither in iPS11 nor in niPS11 cells autophagic turnover of LC3 or p62 were significantly induced (Figure 6A and 6B). Of note, this was also the case for the cancer cell line HCT116. All three cell models remained responsive to bafilomycin A_1_ (indicating basal autophagy) and to starvation, as indicated by an EBSS-dependent reduction of p62. Taken together, these observations lead to the conclusion that the genotoxins BPDE and etoposide do not mount a canonical autophagic response in iPSCs or NPCs.

**Figure 6:**
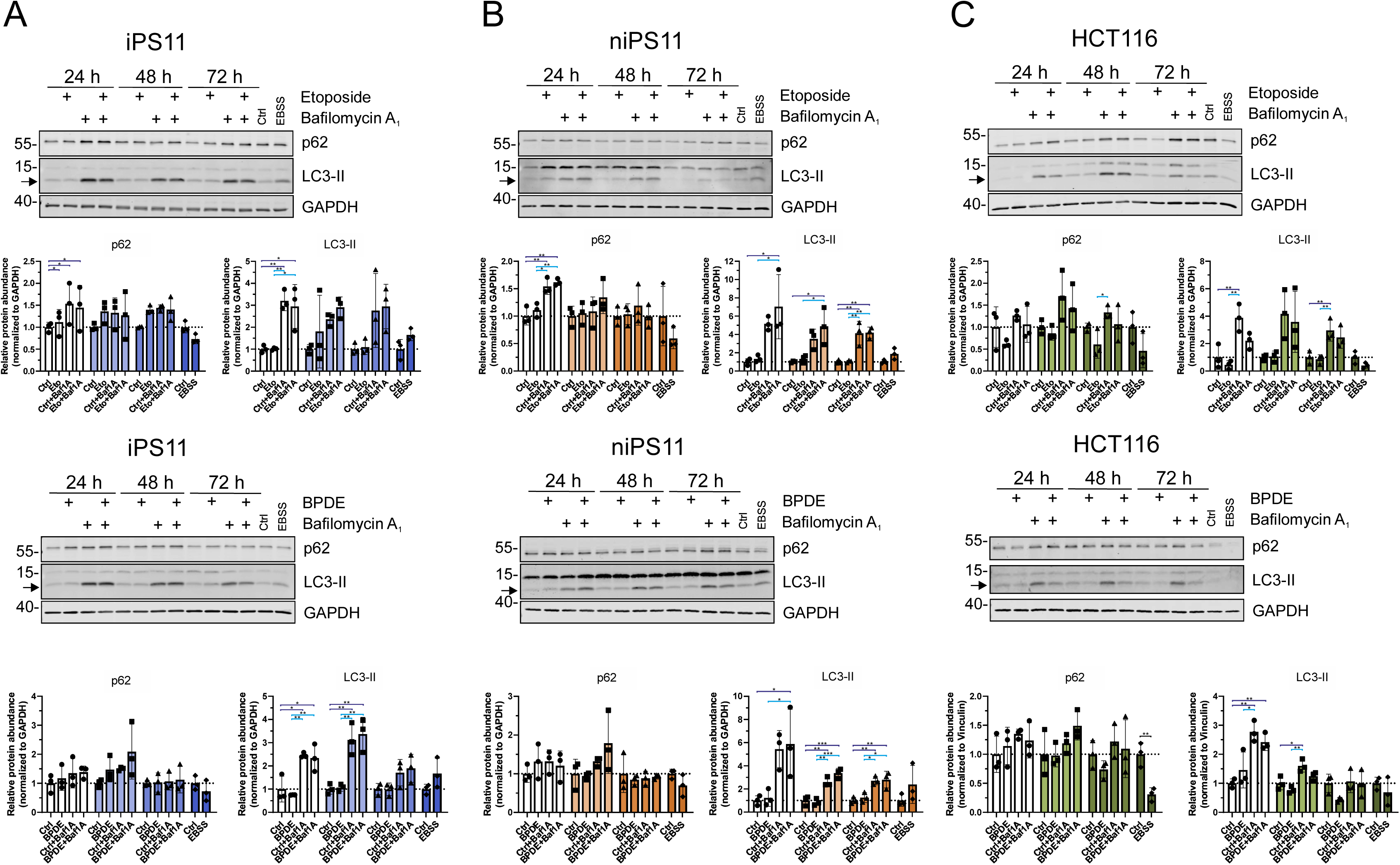
The autophagic flux is not affected by genotoxic treatment. (**A**) iPS11, (**B**) niPS11 and (**C**) HCT116 were treated with corresponding IC_20_ dose for 24, 48 and 72 h. Cells were lysed, and cellular lysates were immunoblotted for SQSTM1/p62, LC3 and GAPDH. One representative immunoblot is shown. Results show mean + SD of three independent experiments. For statistical analysis ordinary one-way ANOVA (Tukeýs multiple comparisons test) and student’s t-test were utilized to compare means of genotoxin-treated samples to DMSO. Significance bars are highlighted in different colors accordingly to involved condition: dark blue: control, light blue: genotoxin; orange: control + bafilomycin A_1_ and genotoxin + bafilomycin A_1_. * p < 0.05, ** p < 0.01, *** p < 0.001, **** p < 0.0001.

### Genotoxin treatment of iPSCs and NPCs does not result in altered expression profiles of autophagy-relevant proteins

In order to get a global overview of the genotoxin-mediated alterations of the cellular proteome, we performed mass spectrometry. In iPS11, proteome changes with regard to biological relevance and statistical significance remained minimal for both treatments (Figure 7A). This was also true for a list of autophagy-relevant genes (orange in Figure 7A). One of the slightly enriched proteins upon BPDE treatment was the transcription factor nuclear ubiquitous casein kinase and cyclin-dependent kinase substrate 1 (NUCKS1), which has been implicated in the regulation of both S phase entry and double-strand break repair (De Angelis et al., 2018; Hume et al., 2021; Maranon et al., 2020; Parplys et al., 2015; Yue et al., 2016). This upregulation was also confirmed by immunoblotting of the samples analyzed by mass spectrometry (Figure 7B). Similar to iPS11, the proteome alterations with regard to autophagy-relevant proteins remained at low levels in niPS11 (possibly except for a downregulation of PRKAR2A upon BPDE treatment and an upregulation of SESN2 upon etoposide treatment). In niPS11, specifically proteins involved in the regulation of mitosis appeared to be upregulated (Figure 7C). Again, this was confirmed by immunoblotting for the candidate proteins polo like kinase 1 (PLK1) and aurora kinase A (AURKA) (Figure 7D). Interestingly, for BPDE treatment, an AURKA downregulation was observed. We next aimed at determining whether these observed changes in protein abundance translated into mitotic alterations. For that we performed immunoblotting (acetylated tubulin), mitotic index assays, and growth curves (Figures 7D and 7E). During mitosis, microtubules become acetylated, and this post-translational modification is important for proper spindle function and chromosome segregation (Piperno et al., 1987; Rasamizafy et al., 2021). We observed increased levels of acetylated tubulin upon BPDE treatment, but this was not the case for etoposide. Phosphorylation of histone H3 at Ser10 is linked to chromosome condensation during mitosis (Hendzel et al., 1997; Wei et al., 1998; Wei et al., 1999). However, we did not detect any differences of H3 Ser10 phosphorylation upon treatment with genotoxins. Finally, proliferation rates appeared to be similar between untreated and treated niPS11 (Figure 7E).

**Figure 7:**
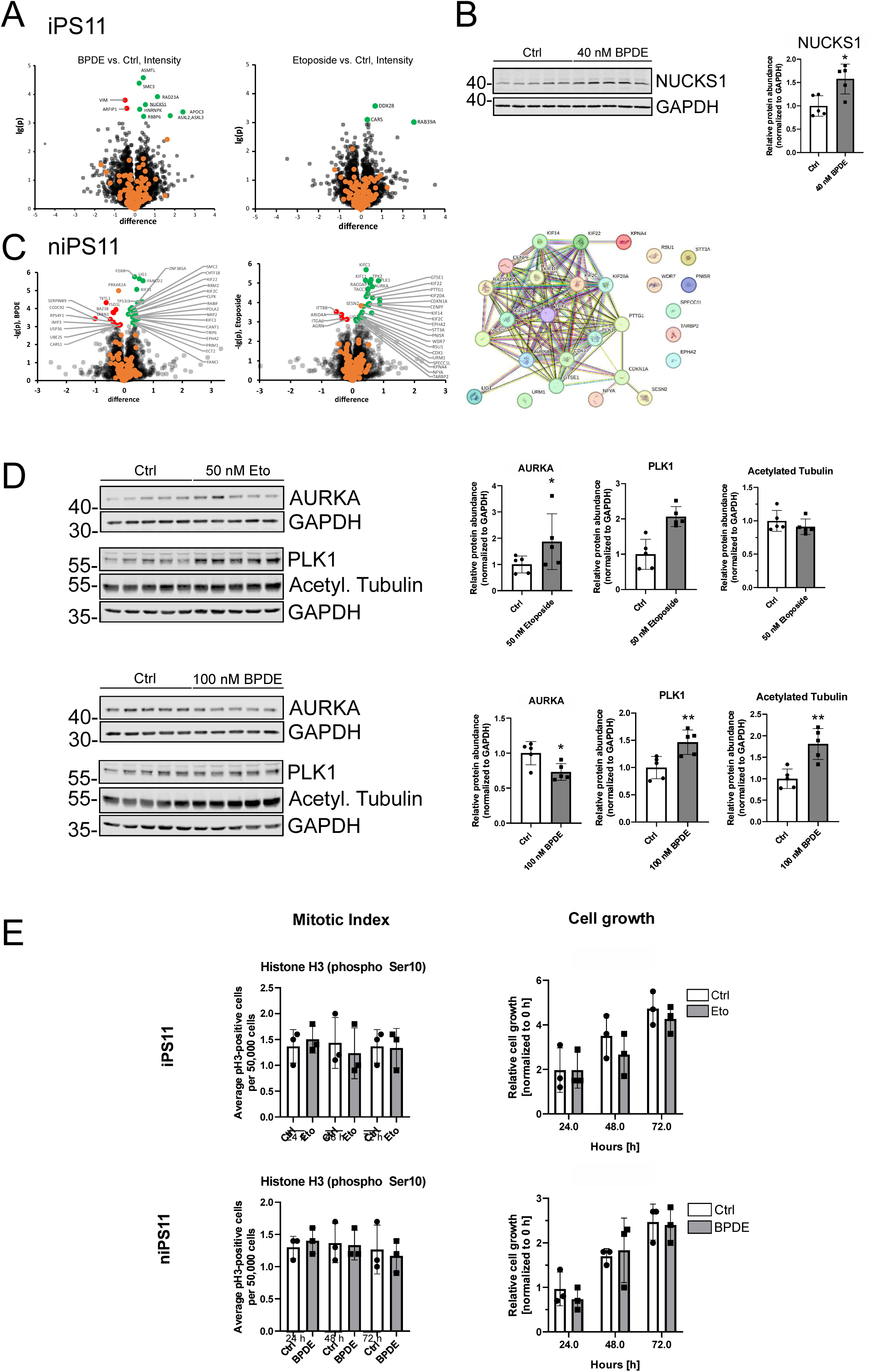
Differential Proteome Analysis of BPDE- and etoposide-treated iPS11 and niPS11. (**A**) Volcano plots based on intensity values for MS-based proteomics of BPDE- or etoposide-treated iPS11. Autophagy-relevant proteins are indicated by orange data points. Proteins with a -lg(p-value) >= 3 significance cutoff are displayed as red (down-regulated) or green (up-regulated) data points. (**B**) Samples from differential proteome analysis were prepared with sample buffer and cellular lysates were immunoblotted for NUCKS1 and GAPDH. The blot with all samples is shown. Results show mean + SD of five independent experiments. For statistical analysis student’s t-test were utilized to compare means of genotoxin-treated samples to DMSO. (**C**) Volcano plots for MS-based proteomics of BPDE- or etoposide-treated niPS11. Autophagy-relevant proteins are indicated by orange data points. Proteins with a -lg(p-value) >= 3 significance cutoff are displayed as red (down-regulated) or green (up-regulated) data points. Proteins upregulated by etoposide were submitted to a functional enrichment analysis using STRING (v12.0, https://string-db.org) yielding the displayed protein-protein interaction network, for which the gene ontology term GO:190304 (Mitotic cell cycle process) was most prominent. (**D**) Samples from differential proteome analysis were prepared with sample buffer and cellular lysates were immunoblotted for Aurora kinase A (AURKA), polo like kinase 1 (PLK1), acetylated tubulin (K40), and GAPDH as loading control. The blot with all samples is shown. Results show mean + SD of five independent experiments. For statistical analysis student’s t-test were utilized to compare means of genotoxin-treated samples to DMSO. (**E**) For mitotic index assay, niPS11 were treated with 50 nM etoposide and 100 nM BPDE for 24, 48 and 72 h. Cells were fixed, permeabilized and stained for phospho-H3 Ser10 and DNA marker DRAQ7 and were analyzed by flow cytometry. 50,000 cells per experiment were quantified and results show mean + SD of three independent experiments. For statistical analysis student’s t-test was utilized to compare means of genotoxin-treated samples to DMSO. Growth curve analysis was performed by Trypan Blue Assay. niPS11 were treated with 50 nM etoposide and 100 nM BPDE for 24, 48 and 72 h. Cells were detached and stained by Trypan Blue Stain to count the number of living cells. Results show mean + SD of three independent experiments. For statistical analysis student’s t-test were utilized to compare means of genotoxin-treated samples to DMSO. * p < 0.05, ** p < 0.01, *** p < 0.001, **** p < 0.0001.

In summary, our data indicate that the environmental genotoxin BPDE and the topoisomerase II inhibitor etoposide do not activate the autophagic program in iPSCs and NPCs. Although niPS11 show a slight activation of ULK1 upon treatment, this does not result in increased autophagic flux. Generally, both cell types mount an autophagic response upon starvation, indicating that a functional autophagy machinery is present in these cells.

## DISCUSSION

Several previous works indicate that DNA-damaging drugs induce an autophagic response (Bordin et al., 2013; Eliopoulos et al., 2016; Galati et al., 2019; Juretschke & Beli, 2021; Katayama et al., 2007; Rodriguez-Rocha et al., 2011; Vanzo et al., 2020). However, the vast majority of these analyses has been performed in cancer cell models, and the effect of DNA damage on stem cells remains largely unknown. Here, we aimed at investigating how induced pluripotent stem cells and thereof differentiated neural progenitor cells react to genotoxin treatment with regard to the autophagy signaling pathway. We observed that both iPSCs and NPCs are generally capable of mounting a strong starvation-induced autophagy, confirming the functionality of the autophagy machinery in these cell models. However, the environmental genotoxin BPDE and the topoisomerase II inhibitor etoposide do not induce a canonical autophagy response. We observed that these compounds only moderately alter the global cellular proteome in iPSCs. In NPCs, mitosis-regulatory proteins were differentially expressed, but this does not result in changes of the mitotic index or the proliferation rates.

Autophagy generally represents a cytoprotective stress response. It has been demonstrated by several groups that DNA damage-inducing agents or treatments induce an autophagic response. Katayama et al. reported that temozolomide and etoposide induced an autophagy-dependent increase in ATP production in multiple glioma cell lines (Katayama et al., 2007). Autophagy induction has also been reported for several other DNA-damaging compounds or treatments (Bordin et al., 2013; Galati et al., 2019; Vanzo et al., 2020). The differences between these reports and our data might be caused by different cell models. In all mentioned works, cancer cell lines were used. However, we also do not observe a significant autophagic response in HCT116 upon BPDE or etoposide treatment. Here, additional differential parameters such as duration of treatment or concentration of compounds need to be considered. We have deliberately chosen IC_20_ concentrations for our studies in order to avoid too extensive cell death and to enable subsequent further differentiation of stem cells. Although we confirmed that these concentrations led to significant DNA damage, we cannot exclude that higher concentrations would result in a more pronounced autophagy activation. With regard to the treatment, it has been previously reported that radiation-induced DNA damage induces autophagy in HCT116 (Alotaibi et al., 2016; Qased et al., 2013). However, the extent and persistence of DNA damage might again differ between the different treatments. In several reports it has been described that autophagy inhibition enhances the cytotoxic effects of DNA-damaging chemotherapy. However, a sensitization of cells to genotoxic drugs by pharmacological inhibition of the autophagic pathway does not necessarily imply that the genotoxic drugs themselves directly induce autophagy. In other words, autophagy might represent a cytoprotective “bystander” effect that rather acts as a counter-mechanism to cell death induction. In this case, non-lethal concentrations of a DNA-damaging drug might not be sufficient to elicit an autophagy response.

A further level of complexity arises from the large number of canonical and non-canonical autophagy signaling pathways. Park et al could demonstrate that chaperone-mediated autophagy (CMA) is upregulated in response to DNA damage and mediates the regulated degradation of CHK1 (Park et al., 2015). Notably, they also reported that cells were more sensitive to several genotoxins (methylmethanesulfonate, cisplatin, paraquat, hydroxyurea, etoposide, camptothecin) when CMA was blocked, whereas blockade of canonical autophagy sensitized cells only to alkylating agents (methylmethanesulfonate and cisplatin) (Park et al., 2015). Accordingly, it might be worthwhile to analyze CMA and/or alkylating agents in our cellular model systems. Another autophagic signaling pathway that has been associated with DNA damage is Golgi membrane-associated degradation (GOMED) (Sakurai et al., 2023). This pathway requires 1) ULK1 and 2) dephosphorylation of ULK1 at Ser638 (Sakurai et al., 2023; Torii et al., 2016). We do not observe a significant alteration of ULK1 Ser638 phosphorylation status—at least in iPSCs—we currently do not think that GOMED plays a major role in this cellular system.

One central aspect of our study was the usage of iPSCs and NPCs and to analyze DNA damage-induced autophagy in stem cells. Generally, autophagy is supposed to provide energy in order to maintain both cell cycle arrest and DNA repair activities. Possibly, this is not desired in stem cells, as the genome is “too valuable”. Accordingly, cytoprotective stress responses are not preferred over cell death mechanisms, since for a whole organism it is beneficial that single damaged cells become depleted and are replenished by non-harmed cells. In contrast, tumor cells do not have to be so stringent with regard to their genomic integrity. In this regard, it would be interesting to investigate induction of autophagy in neuronal cells differentiated from iPSCs and NPCs, since they are rather post-mitotic and likely rely on adaptive stress responses in order to avoid undesired cell loss.

Our proteome analysis revealed that mitosis-regulatory factors are upregulated in NPCs upon treatment with genotoxic compounds. We speculate that these alterations might represent an attempt to counteract the G2/M arrest caused by the genotoxins. Ultimately, we do not observe differences in the mitotic index or the proliferation rate. Even these “adjustments” of mitosis-regulating factors were not observed in iPSCs, again indicating that the stem cell pool is strictly controlled with regard to genomic integrity and mitosis.

On the molecular level, a direct crosstalk between DNA damage response factors and the autophagy signaling machinery is well established. Our analyses so far have addressed early (ULK1 activation status) and late autophagic events (LC3 and p62 turnover). However, the direct crosstalk between DNA damage-sensing factors and autophagy initiation in our cellular model systems awaits further clarification. We observe that treatment with both genotoxins induces phosphorylation of p53 and H2AX, respectively, indicating that the DDR-inducing kinases become activated. However, this does apparently not translate into a sustained autophagy activation. With regard to transcriptional control of the autophagic response via p53 or other factors (e.g., TFEB or FOXK), we at least do not observe any differences on the proteomic level. However, we have not yet analyzed alterations on the transcriptional level. Meira de Amorim et al. recently reported that BPDE exposure enhances gene expression of cell cycle arrest related genes, but the authors did observe an impact on the cell cycle (Meira de Amorim et al., 2024). Specific alterations of autophagy-related genes were not reported. Future analyses will also focus on the ATM/ATR/DNA-PK-dependent control of the autophagy regulators AMPK and mTOR, respectively. At the moment we speculate that—although a DNA damage response is initiated—the signaling cascade towards autophagy is blocked at an early stage. Notably, we detect an upregulation of SESN2 in NPCs upon etoposide treatment, a protein that has been shown to regulate autophagy via mTOR, AMPK, and ULK1 (Lu et al., 2023). As we do not observe significant ULK1 activation upon genotoxin treatment, future studies will need to address if inhibition of mTOR or activation of AMPK are affected, or if the blockade might occur on the level of the interaction between SESN2 and ULK1.

In summary, it appears that neither the environmental toxin BPDE nor the topoisomerase II inhibitor etoposide elicit a significant autophagic response in iPSCs or thereof differentiated NPCs, although these cell models mount a “regular” autophagic response upon starvation. These observations indicate that stem and progenitor cells do not tolerate adaptive cellular stress responses if genomic stability and integrity is endangered. Future studies need to address 1) how an autophagic response to genotoxic stress is suppressed and 2) whether alternative or non-canonical forms of autophagy are executed instead.

## MATERIAL AND METHODS

### Antibodies and reagents

Antibodies against ULK1 (Cell Signaling Technology, Danvers, MA, USA, #8054, 1:1000), phospho ULK1 Serine 757 (Cell Signaling Technology, Danvers, MA, USA, #6888, 1:1000), phospho ULK1 Serine 638 (Cell Signaling Technology, Danvers, MA, USA, #14205, 1:1000), GAPDH (abcam, Cambridge, UK, #ab8245, 1:5000), LC3B (MBL, Woburn, MA, USA, #M-152–3, 1:200 for IF and Cell Signaling Technology, Danvers, MA, USA, #2775, 1:1000 for WB), SQSTM1/p62 (PROGEN, Heidelberg, Germany, #GP62-C, 1:1000), Vinculin (Sigma-Aldrich, St. Louis, MO, USA, #V9131, 1:2000), XPC (Cell Signaling Technology, Danvers, MA, USA, #14768, 1:1000), p53 (Cell Signaling Technology, Danvers, MA, USA, #9282, 1:1000), phospho p53 Serine 10 (Cell Signaling Technology, Danvers, MA, USA, #9284, 1:1000), OCT4a (Cell Signaling Technology, Danvers, MA, USA, #2840, 1:1000), Nestin (Merck Millipore, Darmstadt, Germany #MAB5326, 1:200), PAX6 (Biolegend, San Diego, CA, USA, #901302, for WB: 1:1000, for IF: 1:100) , NUCKS1 (Proteintech, Chicago, IL, USA, #12023-2-AP, 1:1000), γH2AX (For WB: Cell Signaling Technology, Danvers, MA, USA, #2577, 1:1000 and for IF: Merck Millipore, Darmstadt, Germany #05-636, 1:100), 53BP1 (Novus bio, Centennial, CO, USA, 1:2500), Aurora A (Cell Signaling Technology, Danvers, MA, USA, #144755, 1:1000), acetylated Tubulin (Sigma-Aldrich, St. Louis, MO, USA, #312701, 1:20000), PLK1 (abcam, Cambridge, UK, #ab189139, 1:1000), phospho Histone 3 Serine 10 (Cell Signaling Technology, Danvers, MA, USA, #3377, 1:1600) were used. For WB, IRDye 800- or IRDye 680-conjugated secondary antibodies were purchased from LI-COR Biosciences (Lincoln, NE, USA, #926-68077, #926-32211 and #926-32210). Secondary antibodies for immunofluorescence analyses and mitotic index assay were purchased from Jackson ImmunoResearch (Alexa Fluor 488-AffiniPure Goat Anti-Mouse IgG, 1:500, #115–545-003; Alexa Fluor 647-AffiniPure Goat Anti-Mouse IgG, 1:500, #115–605-003; Alexa Fluor 647-AffiniPure Goat Anti-Rabbit IgG, 1:500, #111–605-144 and Alexa Fluor 488-AffiniPure Goat Anti-Rabbit IgG, 1:500, #111– 545-003). Other reagents used were bafilomycin A_1_ (Sigma-Aldrich, St. Louis, MO, USA, #B1793), DMSO (PanReac AppliChem, Darmstadt, Germany, #A3672), 70% ethanol (VWR, Radnor, PA, USA, #85825.360), thiazolyl blue tetrazolium bromide (MTT, ROTH, Karlsruhe, Germany, #4022.3), Pierce BCA Protein Assay Kit (Thermo Fisher Scientific, Waltham, MA, USA, #23225), DRAQ7™ (abcam, Cambridge, UK, #ab109202, 1:100), Benzo[a]pyrene diol epoxide (Santa Cruz, Dallas, TX, USA, #sc-503767) and etoposide (abcam, Cambridge, UK, #ab120227).

### Cell lines and cell culture

iPSCs were cultured in mTeSR Plus (Stemcell Technologies, Vancouver, Canada, #100-0276) supplemented with 100 units/mL Penicillin-Streptomycin (P/S; 10,000 U/ml Penicilin, 10,000 µg/ml Streptomycin) (Thermo Fisher Scientific, Waltham, MA, USA, Gibco, #15140122). Neural progenitor cells were differentiated and cultured in self-prepared medium as previously described (Zink et al., 2021). Maintenance medium (sm-) consists of Neuralbasal A medium (Thermo Fisher Scientific, Waltham, MA, USA, #10888022), DMEM/F12 HEPES (Thermo Fisher Scientific, Waltham, MA, USA, #31330038), B27 supplement without vitamin A (Thermo Fisher Scientific, Waltham, MA, USA, #12587010), N2 supplement (Thermo Fisher Scientific, Waltham, MA, USA, #17502048) and L-Glutamine (200 mM) (Thermo Fisher Scientific, Waltham, MA, USA, Gibco, #25030081). Medium was stored up to 2 weeks at 4°C or aliquoted and frozen at -20°C. Before usage, 3 µM CHIR99021 (Cayman Chemical, Ann Habor, MI, USA, #Cay13122-5), 500 nM Purmorphamine (Miltenyi Biotec, Bergisch Gladbach, Germany, #130-104-465) and 150 µM (+)-Sodium L-ascorbate (Vitamin C) (Merck, Sigma-Aldrich, St. Louis, MO, USA, #A4034) were supplemented to the maintenance medium (sm+). Both cell types were cultivated in 6 well plates coated with Geltrex (Thermo Fisher Scientific, Waltham, MA, USA, #a1413302). Coated plates were incubated at 37°C for 1 h before usage. After thawing of iPSCs and NPCs, cells were supplemented with 10 µM Rock inhibitor/Y-27632 (Dihydrochloride) (Stemcell Technologies, #72304) for 24 h. For passaging and seeding, iPSCs were treated with ReLeSR (Stemcell Technologies, Vancouver, Canada, #100-0483) accordingly to manufactureŕs instructions and NPC were passaged by Accutase (Stemcell Technologies, Vancouver, Canada, #07922). HCT116 were cultured in McCoy medium (Thermo Fisher Scientific, Waltham, MA, USA, Gibco, #36600-021) supplemented with 100 units/mL Penicillin-Streptomycin and 10% fetal bovine serum (Sigma-Aldrich, St. Louis, MO, USA, #F9665, LOT 0001655439) and passaged with Trypsin/EDTA 0.05% (Thermo Fisher Scientific, Waltham, MA, USA, Invitrogen, #2530096). All cells were cultured and treated at 37 °C and 5% CO_2_ in a humidified atmosphere.

### Stimulation

To circumvent autophagy induction by starvation due to rapid proliferation of the stem cells the medium was exchanged on a daily basis. Thereby, cells were exposed to etoposide for 24 h and subsequent medium exchanges were without etoposide. In comparison, BPDE was freshly supplemented to the cells every day. On the day of sample harvesting, the medium was exchanged 4 h before lysis and 40 nM bafilomycin A_1_ were supplemented to the cells 2 hours before lysis. For showing autophagic response in the experiments, cells were washed with DPBS and incubated with EBSS (Thermo Fisher Scientific, Waltham, MA, USA, #24010-043) and corresponding medium control for 2 h. A detailed treatment scheme is depicted in supplemental Figure 2.

For the analysis of differential proteome, cells were treated like described above. Thereby, analysis for etoposide treated cells were performed at 24 h and treated with BPDE after 48 h.

### Cell viability assay

For assessment of cytotoxicity the MTT (3-(4,5-dimethylthiazol-2-yl)-2,5-diphenyltetrazolium bromide) assay was performed. Cells were seeded in 96-well plates (density per well: iPS11: 2500 cells; niPS11: 15000 cells; HCT116: 300-1250 cells). 3-4 days after seeding, cells were treated with different concentrations of BPDE or etoposide, 0.1% DMSO as a solvent control for 24, 48 and 72 h. After the incubation time, 20 µL of a 5 mg/mL MTT stock solution (ROTH, Karlsruhe, Germany, #4022.3) were added to the cells and they were incubated at 37 °C and 5% CO_2_ in a humidified atmosphere for 30 min. Upon removal of the MTT-containing medium 100 µL DMSO per well were added for extraction of the formazan. Absorbance was measured at 570 nm and 650 nm (reference) with a microplate reader (SynergyMx, BioTek, Winooski, VT, USA). After subtraction of the reference value, the mean of the absorbance of the solvent control was set as 100% and the relative viability was calculated for each sample.

### Immunoblotting

For SDS PAGE and western blotting, cells were washed with DPBS and lyzed with RIPA buffer (150 mM Sodium cloride, 50 mM Tris-HCl, 1% Nonidet-40, 0.5% Sodium deoxycholate [w/v], 0.1% SDS [w/v] X PhosSTOP [Roche, Basel, Switzerland, #4906837001]), 1X protease inhibitor cocktail [Roche, Basel, Switzerland, #4693132001]) for 20 min on ice and the lysates were cleared by centrifugation at 18000 rcf and 4 °C for 20 min and quickly frozen in liquid nitrogen. Protein concentration was determined by BCA assay and sample buffer was added (62.5 mM Tris, 8.6% [v/v] glycerol, 2% [w/v] SDS, 33.3 µg/mL bromophenol blue, 1% [v/v] β-mercaptoethanol). Samples were heated at 95 °C for 5 min and then equal amounts of protein (25 µg) were subjected to SDS-polyacrylamide gels. After separation by SDS-PAGE, proteins were transferred to PVDF membranes (Merck, Darmstadt, Germany, #IPFL00010), blocked with 5% milk powder in TBST or EveryBlot Blocking Buffer (Bio-Rad, Hercules, CA, USA, #12010020) and analyzed using the indicated primary antibodies followed by appropriate IRDye 800- or IRDye 680-conjugated secondary antibodies (LI-COR Biosciences, Lincoln, NE, USA). Fluorescence signals were detected using an Odyssey Infrared Imaging system (LI-COR Biosciences, Lincoln, NE, USA) and signals were quantified with Image Studio Lite 5.2 (LI-COR Biosciences, Lincoln, NE, USA).

### Immunofluorescence

For immunofluorescence microscopy, cells were seeded on glass coverslips in 24-well plates. Coverslips for iPSCs and NPCs were coated with geltrex and incubated at 37°C for 1 h. After treatment, cells were fixed in 4% paraformaldehyde for 15 min on room temperature, quenched with 50 mM NH_4_Cl for 15 min and permeabilized with either 50 µg/mL digitonin (Sigma-Aldrich, St. Louis, MO, USA, #D141) or 0.5% Triton X-100 for 10 min. Fixed samples were blocked with 3% BSA (Roth, Karlsruhe, Germany, #8076) for 30 min and incubated with primary antibodies diluted in 3% BSA for 1 h at RT. Samples were then washed three times with DPBS, incubated with the appropriate secondary antibodies and 2 µg/mL DAPI (Roth, Karlsruhe, Germany, #6335.1) diluted in 3% BSA for 1 h and washed three times with DPBS. Afterwards, cells were embedded in ProLong Glass Antifade Mountant (Thermo Fisher Scientific, Waltham, MA, USA, #P36980). Images were recorded with an Axio Observer 7 fluorescence microscope (Carl Zeiss Microscopy, Oberkochen, Germany) using a 40x/1,4 Oil DIC M27 Plan-Apochromat objective (Carl Zeiss Microscopy, Oberkochen, Germany) and an ApoTome 2 (Carl Zeiss Microscopy, Oberkochen, Germany).

### Mitotic Index Assay

For determination of the mitotic index, cells were detached and fixed in cold 70% ethanol and stored at 4°C for up a week. During sample preparation, cells were permeabilized in 0.25% Triton X-100 on ice for 15 min and incubated with rabbit anti-H3 phospho-Ser10 antibody in wash buffer, consisting of 1% BSA in DPBS, overnight at 4°C in constant rotation. All samples were washed twice in wash buffer and incubated anti-rabbit Alexa 488 diluted in wash buffer for 30 min at RT in the dark. After a final wash in washing buffer each cell pellet was re-suspended in 500 µL DPBS containing 3 μM Draq7 for DNA staining, filtered through a cell strainer, processed in a BD LSR Fortessa flow cytometer with FACSDIVA software and analyzed with FlowJo software.

### Growth curve analysis

To investigate the impact on proliferation, niPS11 were treated and cultured as described above and detached by using Accutase. Cells were incubated for 5 min at 37°C and subsequently centrifuged at 60 x g for 3 min. Cell pellet was resuspended in 250 µL sm+ medium. Afterwards, 20 µL of cell suspension were mixed with 20 µL Trypan Blue Stain (0.4%) (Thermo Fisher Scientific, Waltham, MA, USA, Gibco, #15250-061) and measured by Luna Automated Cell Counter (Biocat, Heidelberg, Germany, Model #L10001). Number of daily measured cells were divided by the cell number at 0 h.

### Mass Spectrometry (MS)-based Proteomics

#### SAMPLE PREPARATION

Sample preparation was performed as described (Sinatra et al., 2022; Sprengel et al., 2025).

#### LC-MS Analysis

LC-MS analysis was performed essentially as described (Sinatra et al., 2022) using a QExactive Plus Hybrid Quadrupole-Orbitrap mass spectrometer (Thermo Fisher Scientific, software versions: Xcalibur software: version 4.5.474.0, LC: Thermo Scientific SII for Xcalibur 1.7.0.468, MS: Q Exactive Plus - Orbitrap MS 2.12 build 3134), operated in positive mode and coupled with a nano electrospray ionization source connected with an Ultimate 3000 Rapid Separation liquid chromatography system (Dionex/Thermo Fisher Scientific, Idstein, Germany) equipped with an Aurora Ultimate C18 column (75 µm inner diameter, 25 cm length, 1.7 µm particle size from Ion Opticks) as separation column and an Acclaim PepMap 100 C18 column (75 µm inner diameter, 2 cm length, 3 µm particle size from Thermo Fisher Scientific) as trap column, using a 120 min LC gradient. Capillary temperature was set to 250 °C and source voltage to 1.5 kV.

For iPS11 sample set analysis, using a data-dependent acquisition (DDA) top ten method, MS survey scans were carried out over a mass range from 350 to 2000 m/z at a resolution of 140 000. The automatic gain control target was set to 3 000 000, and the maximum fill time was 80 ms. The 10 most intensive peptide ions with charge states +2 and +3 were selected (2 m/z isolation window, 1700 intensity threshold, minimum automatic gain control target 1000), fragmented by high-energy collisional dissociation (normalized collision energy 30), and fragments were analyzed (scan range 200–2000 m/z, resolution 17,500, target for automatic gain control 10,000, maximum injection time 60 ms). Selected precursors were dynamically excluded for 100 s.

For the niPS11 sample set, data-independent acquisition (DIA) was used on the same instrument with otherwise same parameters. One survey scan was followed by six DIA scans, respectively, all with 35,000 resolution and 3,000,000 as target for automatic gain control. Survey scans were carried out over a mass range from 400 to 1650 m/z and the maximum fill time was 200 ms. DIA scans had 200 m/z as fixed first mass and the normalized collision energy set to 30 with automatic maximum injection time, and were performed on 27 isolation windows, each of 20 m/z width, with equidistant centers (19 m/z distance) starting at 410 m/z and ending at 904 m/z.

#### Data Analysis

For the iPS11 sample set, data analysis was performed as described (Sinatra et al., 2022; Sprengel et al., 2025) using MaxQuant (version 2.5.2.0, Max Planck Institute for Biochemistry, Planegg, Germany) and a human sequence database (UniProtKB, downloaded on 12/21/2023, 82685 entries). For the niPS11 sample set, data analysis was performed using DIA-NN (version 1.9.2, (Demichev et al., 2020)) and a human sequence database (UniProtKB, downloaded on 07/08/2024, 82518 entries). For both analyses, methionine oxidation and N-terminal acetylation as well as carbamidomethylation at cysteine residues were considered as variable and fixed modifications, respectively, and a false discovery rate of 1% on protein and peptide levels was set as identification threshold. Statistical analysis was performed as described (Sinatra et al., 2022; Sprengel et al., 2025) but using a -lg(p-value) >= 3 significance cutoff instead of SAM 5% FDR.

### Statistical analysis

All IC_20_ values were calculated with non-linear regression using GraphPad Prism 8.0.2. For quantification of immunoblotting experiments, the signal of each protein band was divided by the average signal of all protein bands of the respective protein and furthermore normalized to the ratio of the loading control. These normalized ratios were divided by the average normalized ratio of the DMSO controls of all biological replicates to calculate fold changes. The background signal for each membrane was subtracted ahead of quantification. All p-values were calculated with ordinary one-way ANOVA (Tukeýs multiple comparison test) and student’s t-test if not indicated otherwise. For immunofluorescence analyses, puncta, nuclei and co-localization were quantified and analyzed using Biovoxxel ImageJ v1.54p. A punta/foci to nuclei ratio was calculated for each image to determine the average number of punta/foci per cell, and were normalized by dividing through the mean dot number of the solvent control. All macros used for quantifications are provided in Supplementary Table xxx. 10 representative images from three biological replicates per experiment were analyzed. For all immunofluorescence analyses, results are shown in scatter plot diagrams visualized as mean with standard deviation and p-values were determined by student’s t-test and are shown in the diagrams. All p-values are shown as *p ≤ 0.05; **p ≤ 0.01; ***p ≤ 0.001; ****p ≤ 0.0001.

## Abbreviations

AMPK: AMP-activated protein kinase
ATG: autophagy-related
AURKA: aurora kinase A
BafA_1_: bafilomycin A_1_
BECN1: beclin 1
DDR: DNA damage response
FIP200: focal adhesion kinase family interacting protein of 200 kDa
IC_50_: half maximal inhibitory concentration
iPSC: induced pluripotent stem cells
(MAP1)LC3: (microtubule-associated proteins 1A/1B) light chain 3
mTOR: mechanistic target of rapamycin
NPC: neural progenitor cell
NUCKS1: nuclear ubiquitous casein kinase and cyclin-dependent kinase substrate 1
PLK1: polo like kinase 1
RB1CC1: retinoblastoma 1 inducible coiled-coil 1
SQSTM1: sequestosome 1
V-ATPase: vacuolar-type H^+^-ATPase

## ACKNOWLEDGMENTS

We thank Gerhard Fritz and all members of the research training group GRK2578 for many insightful comments and suggestions.

## AUTHOR CONTRIBUTION STATEMENT

S.A. designed the experiments, performed viability assays, immunoblot analyses, fluorescence microscopy, growth curve and flow cytometry analyses. A.Z. and A.P. gave technical and theoretical support for the cultivation and/or differentiation of iPSCs and NPCs. T.L. and K.S. performed and analysed mass spectrometry experiments. K.S.K., S.W. and M.J.M. gave technical and theoretical support. S.A., T.L., and B.S. analyzed the data and wrote the manuscript. B.S. supervised the project. All authors discussed the results and commented on the manuscript.

## RESOURCE AVAILABILITY

### Lead contact

Further information and requests for resources and reagents should be directed to and will be fulfilled by the Lead Contact, Björn Stork.

