## Supplemental Information for "Canonical autophagy remains inactive in induced pluripotent stem cells and neuronal progenitor cells following DNA damage induced by BPDE or etoposide"


**AUTHORS/AFFILIATIONS**

Seda Akgün[1], Thomas Lenz[2], Annika Zink[3], Karina Stephanie Krings[1], Sebastian Wesselborg[1], María José Mendiburo[1], Alessandro Prigione[3], Kai Stühler[1,2], Björn Stork[1,*]

[1]*Institute of Molecular Medicine I, Medical Faculty and University Hospital Düsseldorf, Heinrich Heine University, 40225 Düsseldorf, Germany*

[2]*Molecular Proteomics Laboratory, Biological Medical Research Center, Heinrich Heine University Düsseldorf, 40225 Düsseldorf, Germany*

[3]*Department of General Pediatrics, Neonatology and Pediatric Cardiology, Medical Faculty and University Hospital Düsseldorf, Heinrich Heine University, 40225 Düsseldorf, Germany*

**CONTACT**

*Corresponding author:

Björn Stork, Universitätsstr. 1, Building 22.03, 40225 Düsseldorf, Germany

**Figure S1: Definition of $IC_{20}$ values fo iPSCs, NPC and HCT116.**

To identify a sublethal $IC_{20}$ dose, cells were treated with different concentrations of BPDE and etoposide for 24, 48 and 72 h while etoposide was only supplemented for 24 h and BPDE was given daily to the cells. After treatment, cell viability was measured using a thiazolylblue (MTT) assay. Results are shown as the mean ± SD of 3-5 independent experiments performed in triplicates for each treatment.

**Figure S2: Treatment scheme used in this study.**

Supp Figure 1

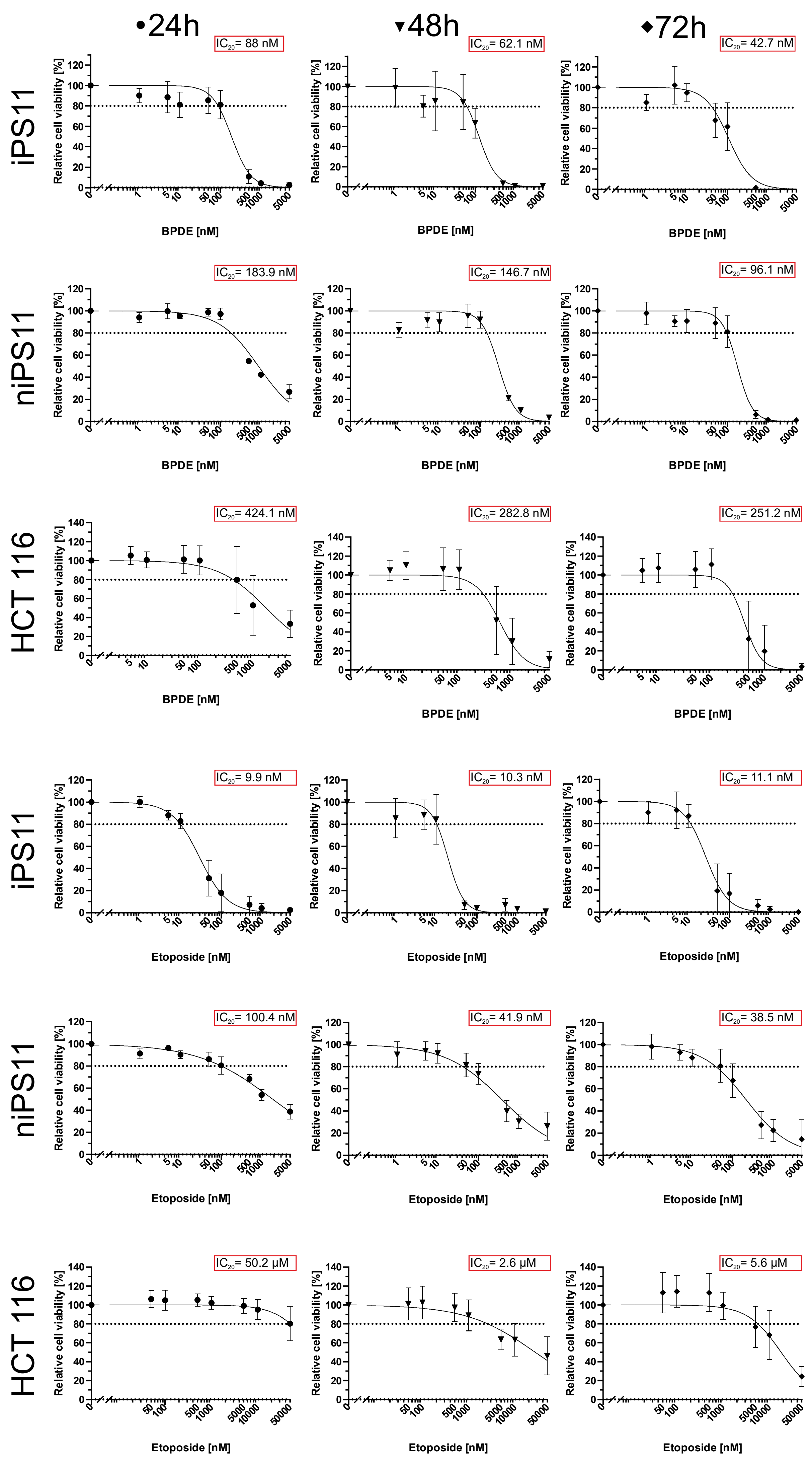

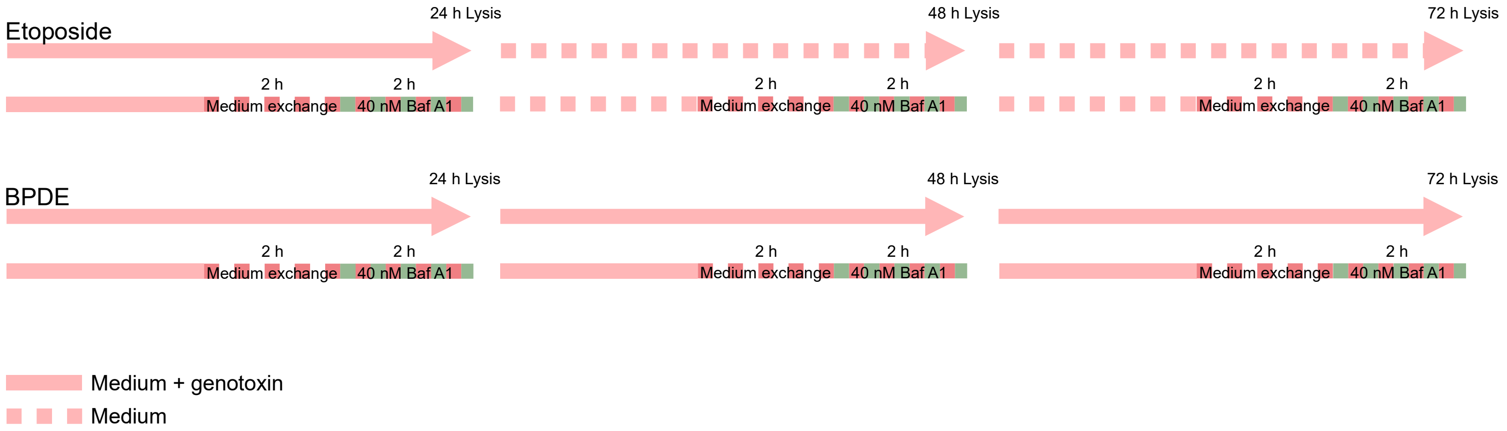


### Supp Figure 2
